# The Lonidamine Derivative H2-Gamendazole Reduces Cyst Formation in Polycystic Kidney Disease

**DOI:** 10.1101/2020.09.09.258160

**Authors:** Shirin V. Sundar, Xia Zhou, Brenda S. Magenheimer, Gail A. Reif, Darren P. Wallace, Gunda I. Georg, Sudhakar R. Jakkaraj, Joseph S. Tash, Alan S.L. Yu, Xiaogang Li, James P. Calvet

## Abstract

Autosomal dominant polycystic kidney disease (ADPKD) is a debilitating renal neoplastic disorder with limited treatment options. It is characterized by the formation of large fluid-filled cysts that develop from kidney tubules through abnormal cell proliferation and cyst-filling fluid secretion driven by cAMP-dependent Cl^−^ secretion. We have examined the effectiveness of the indazole carboxylic acid, H2-gamendazole (H2-GMZ), a derivative of lonidamine, to inhibit these processes and cyst formation using *in vitro* and *in vivo* models of ADPKD. H2-GMZ was effective in rapidly blocking forskolin-induced, Cl^−^-mediated short-circuit currents in human ADPKD cells at 1 μM and it significantly inhibited both cAMP- and EGF-induced proliferation of ADPKD cells with an IC_50_ of 5-10 μM. Western blot analysis of H2-GMZ-treated ADPKD cells showed decreased phosphorylated ERK and hyperphosphorylated Rb levels. H2-GMZ treatment also decreased ErbB2, Akt, and Cdk4, consistent with inhibition of the chaperone Hsp90, and reduced the levels of the CFTR Cl^−^ channel. H2-GMZ-treated ADPKD cultures contained a higher proportion of smaller cells with fewer and smaller lamellipodia and decreased cytoplasmic actin staining, and they were unable to accomplish wound closure even at low H2-GMZ concentrations, consistent with an alteration in the actin cytoskeleton and decreased cell motility. Studies using mouse metanephric organ cultures showed that H2-GMZ inhibited cAMP-stimulated cyst growth and enlargement. *In vivo*, H2-GMZ (20mg/kg) was effective in slowing postnatal cyst formation and kidney enlargement in the *Pkd1*^*flox/flox*^:*Pkhd1-Cre* mouse model. Thus, H2-GMZ treatment decreases Cl^−^ secretion, cell proliferation, cell motility, and cyst growth. These properties, along with its reported low toxicity, suggest that H2-GMZ might be an attractive candidate for treatment of ADPKD.

## INTRODUCTION

Autosomal dominant polycystic kidney disease (ADPKD) affects an estimated 1 in 400-1,000 people worldwide (1-4). ADPKD is characterized by the progressive growth of large fluid-filled cysts in a number of ductal organs, but predominantly in the kidneys. The steady growth and enlargement of kidney cysts ultimately leads to end stage renal disease (ESRD) in about 50% of patients by age 50-60 years and accounts for 6-9% of patients on renal replacement therapy (2,5,6). Many years of productive life are lost due to the debilitating complications of PKD, which include hypertension, hematuria, abdominal pain, and kidney infection (7,8). Blood pressure control has been partially effective in the clinical management of the disease (8,9).

In ADPKD, renal cyst growth requires two mechanisms—proliferation of cyst-lining epithelial cells and secretion of fluid into the cyst lumen (10-12). The cellular second messenger cyclic adenosine monophosphate (cAMP) plays a vital role in promoting both fluid secretion and cell proliferation in ADPKD (13-17). Cyclic AMP acts downstream of G-protein coupled receptors, such as the arginine vasopression (AVP) V2 receptor, to activate PKA. This leads to phosphorylation and activation of CFTR to promote Cl^−^ secretion followed by fluid secretion into the cyst lumen (16,18). Activation of PKA by cAMP also elicits a strong mitogen-activated protein (MAP) kinase response in ADPKD cells, which are primed to be activated by decreased cytosolic calcium caused by PKD mutation (17,19,20). As an understanding of the molecular mechanisms of ADPKD progression has grown in recent years, so has the number of potential therapies (1-3,21-34), including drugs that target cAMP-dependent fluid secretion and target cell growth and proliferation. One such drug is the AVP V2 receptor antagonist, tolvaptan, which has been approved by the U.S. Food and Drug Administration (FDA) for the treatment of adult patients with ADPKD (17,35-39); however, high cost and side effects, including polyuria, nocturia, thirst, and liver complications limit its use in some patients. Many other drugs that have entered clinical trials for ADPKD have been repurposed from non-PKD indications (3,40,41). While many of these drugs show promise, concerns have been raised regarding their long-term use and safety profiles (42-45). To be an effective therapy for ADPKD, drugs will need to be efficacious and well tolerated over decades, since cyst growth is slow. As such, there is a need to develop drugs for PKD that have excellent safety profiles and can target both fluid secretion and cell proliferation to successfully reduce cyst growth over a lifetime while preserving normal renal function.

Indazole-carboxylic acids are a class of drugs with the potential to target the cellular machinery implicated in the pathogenesis of PKD. Lonidamine (LND) is best known for its anti-spermatogenic and anti-cancer properties (46,47). LND has been shown to inhibit cAMP-induced CFTR-mediated anion secretion (48,49) as an open channel inhibitor of CFTR (50), making it and this class of drugs potential candidates for inhibiting cyst-filling fluid secretion in PKD. LND also affects a number of other cellular processes, including lactate transport and the mitochondrial metabolic pathway for aerobic glycolysis and oxygen consumption (51), inducing apoptosis. More recently, one of the derivatives of LND, gamendazole (GMZ), which was developed at the University of Kansas, was shown to inhibit the Hsp90 pathway, although the mechanism is still a matter of investigation (52-54), and to decrease the levels of the proteins Her2 (ErbB2) and Akt in MCF-7 cells (52). In addition, GMZ was found to associate with the eukaryotic elongation factor eEF1A1. GMZ and another derivative H2-GMZ have been under investigation as male contraceptives. As such, they have excellent safety profiles and bioavailability, and their effects on the testis are reported to be reversible (55).

H2-GMZ has been shown (using proteolytic fingerprinting and affinity chromatography) to modulate Hsp90 function by a mechanism similar to the natural product celastrol (53). The molecular chaperone Hsp90 has proven to be an important drug target for the treatment of various neoplastic disorders. Hsp90 is a ubiquitous protein, abundant in the cell and required for normal function (56,57). Hsp90 promotes the maturation and proper folding of more than a hundred substrate or “client” proteins in the cell. Hsp90 inhibitors are of growing therapeutic interest because client proteins are frequently mutated, over-expressed, or functionally active in cancer, thus making these cells more dependent on a finite supply of Hsp90. Various classes of Hsp90 inhibitors currently being tested as anti-cancer agents have been reviewed extensively (58-63). These agents bind distinct Hsp90 domains or co-chaperones such as cdc37, inhibit their function and cause proteasomal degradation of their client proteins, including numerous protein kinases, transcription factors and cell surface receptors, leading to decreased cell proliferation. Many Hsp90 client proteins have been implicated in the pathogenesis of PKD, including but not limited to ErbB2 (64,65), CFTR (66,67), Akt (68,69), cyclin dependent kinases (70), and proteins of the MEK-ERK pathway (71-74). The Hsp90 inhibitors STA-2842 and STA-9090 (ganetespib) have been shown to reduce cyst size and disease progression in mouse models PKD (75-77). However, the role of the Hsp90 chaperone complex in human ADPKD cells has not been investigated.

Ultimately, we are interested in developing H2-GMZ and its derivatives as a therapeutic for ADPKD based on the ability of the parent compound, LND, to block CFTR channel activity and inhibit cell proliferation. As such, we tested whether H2-GMZ can target both fluid secretion and cell proliferation in human ADPKD cells. To this end, we investigated the effect of H2-GMZ on CFTR-mediated Cl^−^ secretion in ADPKD cells and whether H2-GMZ can inhibit Hsp90 client proteins and cell motility. We further investigated the potential of H2-GMZ to target cyst formation and enlargement in a mouse metanephric organ culture model and *in vivo* in a *Pkd1* conditional mouse. Our results suggest that H2-GMZ could be effective against ADPKD cyst growth.

## METHODS

### H2-GMZ

H2-gamendazole (3-[1-(2,4-dichlorobenzyl)-6-trifluoromethyl-1H-indazol-3-yl]-propionic acid) (JWS-2-72) was initially described in (52) and U.S. Patent US8362031B2.

### Cell culture

Primary cultures of human ADPKD cyst-lining epithelial cells and normal human kidney (NHK) cells were obtained from the PKD Biomarkers, Biomaterials, and Cellular Models Core in the Kansas PKD Center. The use of discarded clinical specimens is considered to be not human subjects research by regulatory agencies and the institutional board at the University of Kansas Medical Center. The generation and use of primary ADPKD and NHK cells have been described in detail (78,79). Cells were maintained in a 37^°^C humidified CO_2_ incubator in DMEM/F12 medium containing 1% FBS and supplemented with insulin, transferrin and selenium (ITS). The mouse cortical collecting duct cell line M-1 (80) was maintained in DMEM/F12 medium containing 5% FBS.

### Chloride ion secretion assay

Anion secretion was determined by measuring short-circuit current across human ADPKD cells. Briefly, confluent monolayers of ADPKD cells on Snapwell supports were inserted into modified Ussing chambers (Harvard Apparatus), and both surfaces were bathed in a HCO_3_-Ringer’s solution maintained at 37°C and equilibrated in 5% CO_2_-95% O_2_. Electrodes were positioned within the chambers and short-circuit current (I_SC_) was measured using two dual-voltage-clamp devices (Warner Instruments). Positive I_SC_ reflects the sum of active transport of cations (e.g., Na^+^) from the apical to basolateral surface and anions (e.g., Cl^−^) from the basolateral to apical surface. Benzamil was added to inhibit cation transport via the epithelial Na^+^ channel ENaC. Forskolin, a potent cAMP agonist, was added to stimulate the Cl^−^ current. I_SC_was continuously monitored and recorded with LabChart7 (AD Instruments).

### Cell proliferation assay

ADPKD cell proliferation was measured using the Promega Cell Titer 96 MTT assay kit as described previously (12,13). Cells were plated at a density of 4 × 10^3^ per well in a 96-well cell culture plate in medium containing 1% FBS + ITS. Cells were serum-starved for 24 h and treated either with 100 µM cAMP or 25 ng/ml EGF to stimulate cell proliferation or were placed directly in 5% FBS-containing medium. Increasing concentrations of H2-GMZ or LND were added to the medium to determine their effect on cell proliferation. After 72 h of treatment, MTT assays were performed by adding dye solution to the cells and stopping the reaction 4 h later. Optical densities measured using a spectrophotometer were considered to be directly proportional to the number of viable cells present.

### Mitotic index

ADPKD cells (5 × 10^3^ per well) were plated in 4-well chamber slides in medium containing 1% FBS + ITS. After 24 hours, medium was changed to include 5% FBS. Cells were treated with increasing concentrations of H2-GMZ for 16 h, fixed with 4% paraformaldehyde and permeabilized with 0.2% Triton-X 100. Cells were mounted using ProLong Gold antifade reagent containing DAPI (Thermo Fisher Scientific) and observed by fluorescence microscopy. Eight to 10 separate fields were examined at 10x magnification and the number of mitotic nuclei were counted and expressed as a percentage of the total number of nuclei per field.

### Western blotting

ADPKD cells were seeded on 6-well culture plates and treated with 50 μM H2-GMZ for various time points. Cytoplasmic protein extracts were prepared by lysing cells in ice-cold lysis buffer (10 mM Tris-Cl pH 7.5, 150 mM NaCl, 2 mM EDTA pH 8.0, 1% Triton X-100, 0.5% NP-40, 25 mM glycerol 2-phosphate, 1 mM sodium orthovanadate, 1 mM phenylmethyl sulfonyl fluoride, and 0.1% v/v Sigma protease inhibitor cocktail). Nuclei and other Triton-insoluble components were removed by high speed centrifugation. Protein concentration was measured using the Pierce BCA assay kit. 20 μg total protein was boiled with SDS sample buffer and fractionated on 7.5, 10 or 12.5% SDS-PAGE gels. Proteins were transferred to PVDF membranes and non-specific binding blocked with 5% powdered milk in TBS-T (10 mM Tris-Cl pH 7.5, 150 mM NaCl, and 0.1% Tween 20) for 1 h at room temperature. Blocked membranes were incubated with primary antibodies in 5% powdered milk or 5% BSA (for phospho-proteins) in TBS-T overnight at 4° C. Membranes were then washed three times with TBS-T and incubated with alkaline phosphatase-conjugated secondary antibodies (Sigma) in 5% milk in TBS-T for 30 min at room temperature. The membranes were washed three times with TBS-T and protein bands were visualized using the CDP-star detection reagent (GE Healthcare). Intensity was detected and quantitatively analyzed by the Fluor-S MAX multi-imager system (Bio-Rad). Antibodies used for Western blotting were p-ERK, Actin (Sigma), ERK1, ERK2, ErbB2, Rb, Cdk4 (Santa Cruz), Hsp90, Hsp70, Akt, GAPDH (Cell Signaling), and CFTR (R&D Systems).

### Immunoprecipitation

ADPKD cells were plated at a density of 1 × 10^6^ cells per flask in medium containing 5% FBS. Cells were lysed after 24 h in gentle lysis buffer (10 mM Tris-Cl pH 7.5, 100 mM NaCl, 2 mM EDTA pH 8.0, 1% NP-40, 20 mM sodium molybdate plus protease inhibitors as above) to prevent dissociation of Hsp90 from client proteins. 500 µg lysate was mixed overnight with 0.5 μg ErbB2 antibody and Protein A/G beads, washed four times with wash buffer (gentle lysis buffer without NaCl) with protease inhibitors and boiled in 2x Laemmli sample buffer. Proteins from 20 µg total cell lysate (input) were analyzed alongside the immunoprecipitate by Western blotting.

### Immunofluorescence

ADPKD cells (1 × 10^4^ per well) were plated in 8-well chamber slides, serum starved for 24 h and treated for 48 h with 25 ng/ml EGF. Cells were fixed, permeabilized and blocked with 1% BSA. Cells were then incubated for 1 h with anti-Hsp90 antibody (Cell Signaling), washed with PBS and incubated for 1 h with FITC-conjugated secondary antibody. Slides were mounted with DAPI-containing mounting medium. Images were captured at 60x. Cultured metanephric kidneys were fixed in 4% paraformaldehyde and frozen sections were cut. For staining, sections were subjected to heat retrieval and were incubated with an anti-PCNA antibody (Sigma) and with anti-mouse Texas Red (Jackson Immunoresearch).

### Actin staining

Human ADPKD cells or mouse M-1 cortical collecting duct cells were plated in a 4-well chamber slide in 5% FBS-containing medium. After 24 h, the cells were treated with increasing concentrations of H2-GMZ for 16 h. Cells were fixed in 4% paraformaldehyde and permeabilized using 0.2% Triton-X 100. Cells were then incubated for 20 min with TRITC-conjugated phalloidin, washed with PBS and mounted with DAPI-containing mounting medium. Images were captured at 10x (A) or 60x (B), using the same exposure settings for treated and untreated cells.

### Cell migration assay

Human ADPKD cells were plated on a 12-well cell culture plate in 5% FBS + ITS medium and allowed to form a confluent monolayer. The monolayer was wounded by scratching it with a P-200 pipette tip and the medium was changed to include increasing concentrations of H2-GMZ. Images of three different fields per well were captured at 10x magnification immediately after wounding and at 4, 8 and 24 h post-wounding. The area of the wound was measured at each time point using NIH Image J program. Percentage of wound closure was calculated by subtracting the area of the wound at any given time point from the 0 h wound area and expressing it as a percentage of the 0 h wound area.

### Metanephric organ culture

*Pkd1*^*m1Bei*^ mice were obtained from the Mutant Mouse Regional Resource Center (University of North Carolina, Chapel Hill, NC) and were stabilized onto a C57BL/6 background (>10 backcrosses). This mouse has a point mutation (T to G at 9248 bp) that causes an M to R substitution affecting the first transmembrane domain of polycystin-1 (81). Mouse metanephric kidneys were cultured according to methods described previously (67,82-84). Metanephroi were dissected from embryonic mice and placed on transparent Falcon 0.4 mm cell culture inserts. DMEM/F12 defined culture medium (supplemented with 2 mM L-glutamine, 10 mM HEPES, 5 µg/ml insulin, 5 µg/ml transferrin, 2.8 nM selenium, 25 ng/ml prostaglandin E, 32 pg/ml T3, 250 U/ml penicillin, and 250 µg/ml streptomycin) was added under the culture inserts, and organ cultures were maintained in a 37°C humidified CO_2_ incubator for up to 6 days. 100 µM 8-Br-cAMP was added to induce cyst formation. This was followed by 1-5 µM H2-GMZ or LND for three days. Upon culturing and approximately 24 h later (Day 1) and each day following (Days 2-5), kidneys were photographed using a 2X or 4X objective, and the images were acquired and quantified using the analySIS imaging program (Soft Imaging System). Fractional cyst area was calculated as total tubule dilation area divided by total kidney area.

### Mouse strains and treatments

All animal protocols were approved and conducted in accordance with Laboratory Animal Resources of the University of Kansas Medical Center and Institutional Animal Care and Use Committee regulations. *Pkd1*^*flox/flox*^:*Pkhd1-Cre* mice (73) were generated by breeding *Pkd1*^*flox/+*^:*Pkhd1-Cre* female mice with *Pkd1*^*flox/+*^:*Pkhd1-Cre* male mice. H2-GMZ treatment was carried out on *Pkd1*^*flox/flox*^:*Pkhd1-Cre* mice using daily intraperitoneal (i.p.) injections of 20 mg/kg H2-GMZ from postnatal (PN) day 8 to 18.

## RESULTS

### H2-GMZ inhibits CFTR-mediated Cl^−^ secretion by ADPKD cells

Lonidamine (LND) has been shown to bind directly to CFTR and inhibit CFTR-dependent transepithelial anion currents across rat epididymal cells (48-50). As such, we were interested in testing the effectiveness of the derivative H2-GMZ as an inhibitor of CFTR-mediated Cl^−^ secreted by ADPKD cells. For this, confluent ADPKD cell monolayers were evaluated in Ussing chambers using short-circuit current assays in the presence of forskolin to stimulate Cl^−^ secretion. The epithelial Na^+^ channel ENaC was inhibited using benzamil to eliminate cation absorption. We found that 1 μM H2-GMZ effectively inhibited CFTR-mediated short-circuit currents within minutes (Fig. 1). Increasing concentrations of LND (up to 120 μM) did not have an effect, whereas addition of H2-GMZ after LND treatment, or alone, reduced the anion current significantly (Fig. 1C, D). Thus, H2-GMZ appears to be much more effective than LND in inhibiting CFTR-mediated Cl^−^ secretion across ADPKD monolayers.

**Fig. 1.**
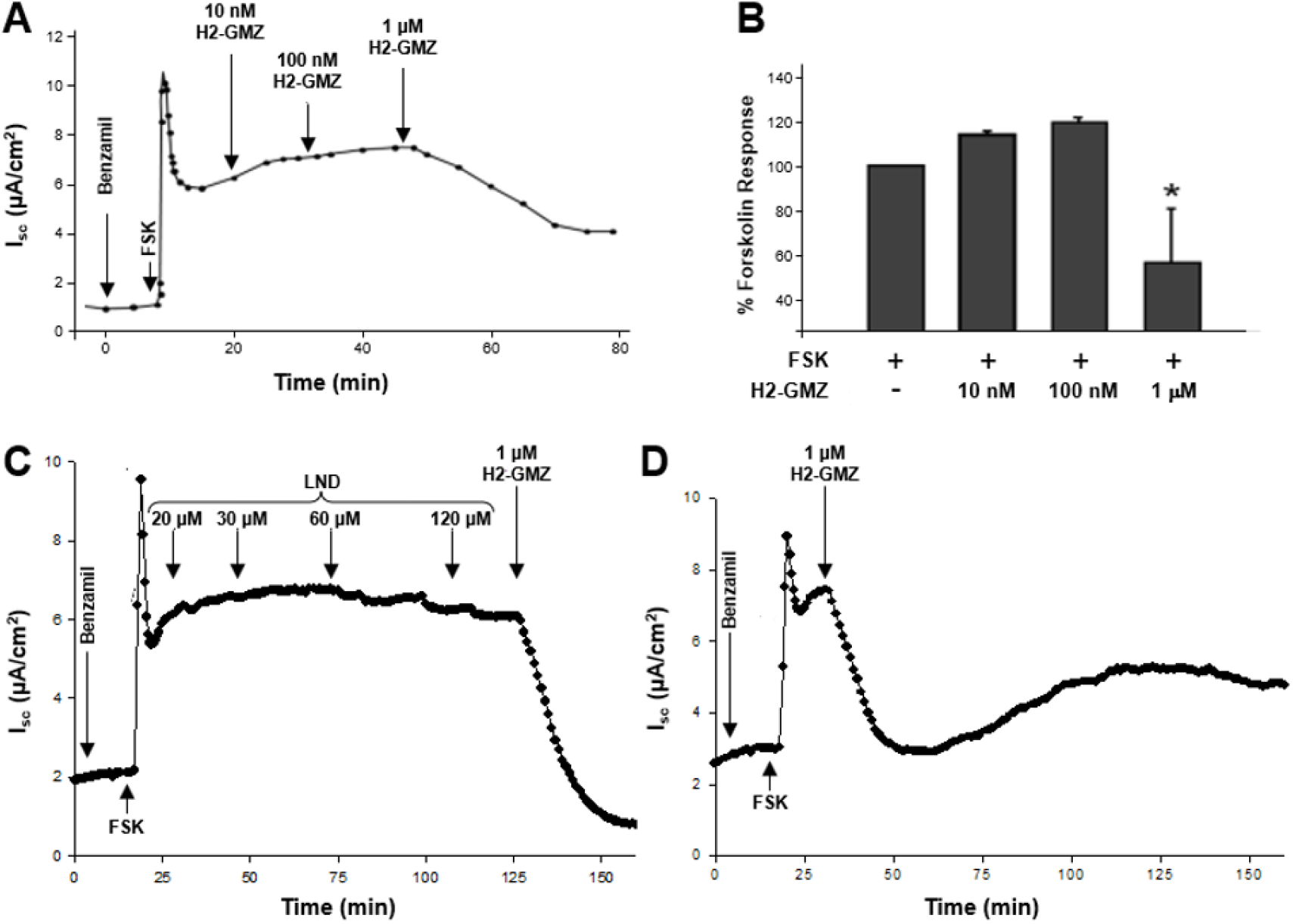
H2-GMZ inhibits CFTR-mediated short-circuit current in ADPKD monolayers. **A** and **B**, Apical treatment with H2-GMZ inhibits forskolin-induced short-circuit current in ADPKD monolayers. Confluent monolayers of human ADPKD cells treated apically with benzamil, then forskolin to first block the ENaC channel and then induce CFTR-current. Increasing concentrations of H2-GMZ were added and the change in current recorded. The graph represents the average of four different monolayers from two ADPKD kidneys. *, the effect was statistically significant at p < 0.05 as determined by ANOVA. **C**, Effect of LND on forskolin-generated current in ADPKD cells followed by 1 μM H2-GMZ, or **D**, with 1 μM H2-GMZ alone.

### H2-GMZ inhibits cell proliferation in ADPKD cells

ADPKD is a neoplastic disorder in which cyst-lining epithelial cells proliferate and secrete fluid into the lumen causing cyst expansion in kidney, liver and other organs. We were interested in examining whether H2-GMZ has an inhibitory effect on the proliferation of ADPKD cells since LND has proven to be an effective anti-proliferative, cancer chemotherapeutic agent. Initial studies with GMZ showed that it inhibits the proliferation of MCF-7 cells with an IC_50_ of ∼100 μM (52). As shown in Fig. 2A, H2-GMZ inhibited the proliferation of ADPKD cells stimulated either with cAMP or EGF and assayed at 72 h. This inhibition occurred in a dose-dependent manner with an IC_50_ of 5-10 μM. In contrast, the IC_50_ was close to 50 μM in the presence of 5% FBS (Fig. 2B). To analyze the effect of H2-GMZ on ADPKD cell proliferation at an earlier time point, we stained cells on chamber slides with the nuclear DAPI stain after treating for 16 h with H2-GMZ. 50 µM H2-GMZ decreased the mitotic index in comparison to untreated controls, and 100 µM H2-GMZ was even more effective (p<0.01, Fig. 2C). The results indicate that H2-GMZ effectively arrests the proliferation of ADPKD cystic cells in culture. The IC_50_ for inhibition of cell proliferation by LND was similar to that for H2-GMZ (Supplemental Fig. 1).

**Fig. 2.**
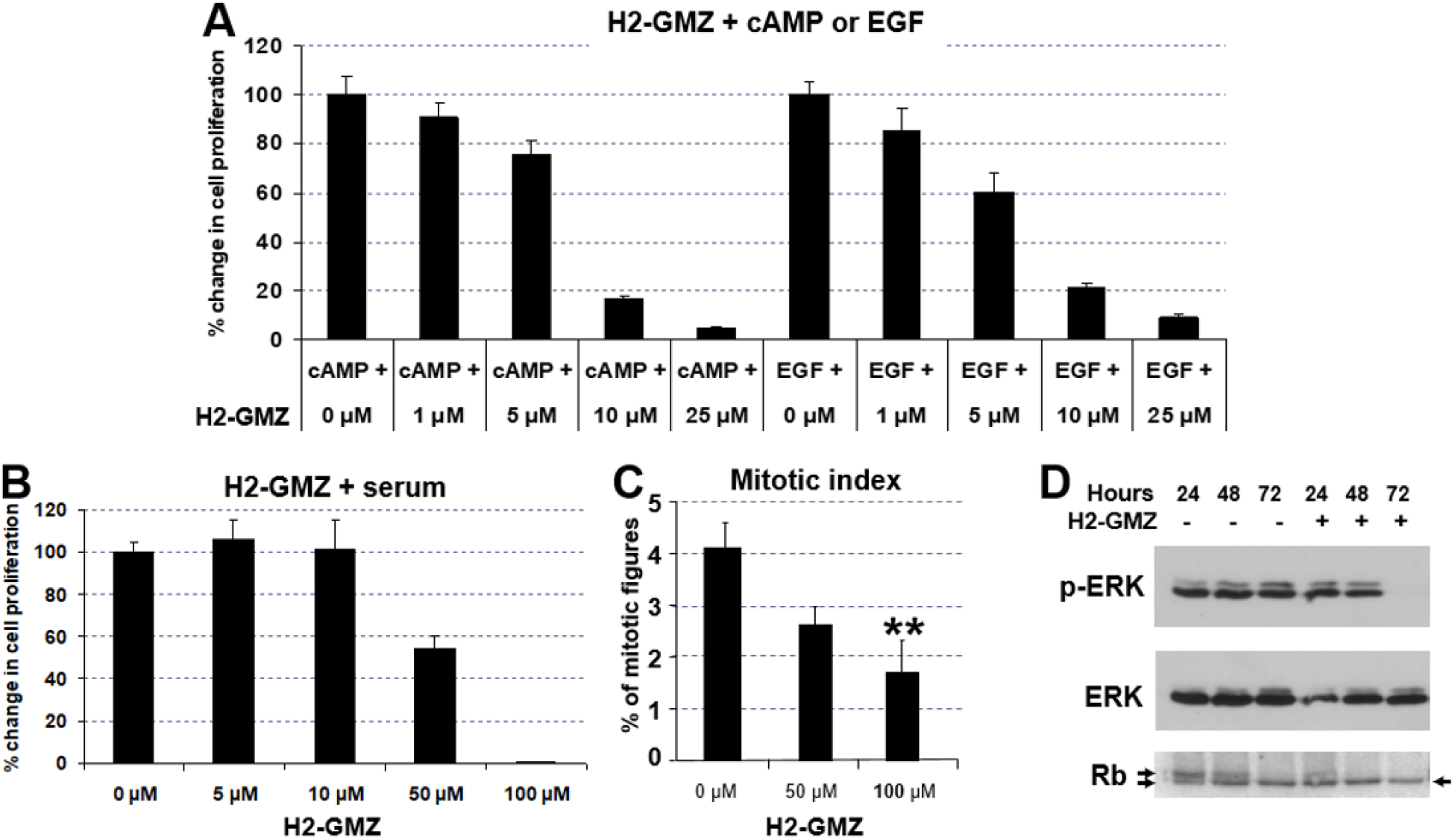
Dose-dependent decrease in the proliferation of primary human ADPKD cells treated with H2-GMZ. **A** and **B**, Dose response of ADPKD cells stimulated with 100 µM cAMP, 25 ng/ml EGF or 5% FBS and treated with H2-GMZ. After 72 hours of treatment, MTT assays were performed. In each group, results were expressed as a percentage of the control group (not treated with H2-GMZ). The error bars indicate a standard error for n = 6. **C**, ADPKD cells stimulated with 5% FBS and treated with H2-GMZ show a significant decrease in the number of mitotic figures. Cells were stained with DAPI to visualize the nuclei. Statistical analysis was done using One-way ANOVA and the Tukey HSD test for pair-wise comparisons. ** indicates p<0.01. **D**, Inhibition of p-ERK and hyperphosphorylated Rb in H2-GMZ treated ADPKD cells correlates with the inhibition of cell proliferation. In growing cells, there is a doublet of hyper- (upper) and hypo- or non-phosphorylated (lower) Rb (retinoblastoma) protein. H2-GMZ treatment decreased the levels of hyperphosphorylated Rb (upper arrow) consistent with decreased cell proliferation. Cytoplasmic extracts were prepared following 24, 48 or 72 hours of 50 μM H2-GMZ treatment in the presence of serum and analyzed by Western blotting.

To determine the mechanism for inhibition of cell proliferation, we tested the effect of H2-GMZ on the levels of phosphorylated ERK (p-ERK). ADPKD cells were treated for 24, 48 or 72 h with 50 μM H2-GMZ. Western blotting with antibodies against p-ERK and total ERK suggested that H2-GMZ-mediated inhibition of cell proliferation occurs at least in part through inhibition of the MEK-ERK pathway (Fig. 2D), which is consistent with previous work that showed that cAMP-driven ERK activation stimulates ADPKD cell proliferation (13,14,19,71) Hyperphosphorylated Rb levels also decreased with H2-GMZ treatment (Fig. 2D, upper band), consistent with inhibition of cell proliferation. Since Rb is an important effector of cell cycle progression, the decrease in hyperphosphorylated Rb as well as p-ERK could contribute to the inhibition of cell proliferation by H2-GMZ.

### H2-GMZ decreases Hsp90 client protein levels in ADPKD cells

Previous studies had shown that GMZ inhibits the Hsp90 pathway in Sertoli cells (52,53). Since the molecular chaperone Hsp90 has been shown to play an important role in maintaining the abnormal proliferative state in various tumor types, we sought to explore the status of Hsp90 in ADPKD cells and whether H2-GMZ inhibited the Hsp90 pathway in these cells. The presence of Hsp90 in ADPKD cells was checked by immunofluorescence, showing localization in the cytoplasm and nucleus (Fig. 3A). The presence of Hsp90 in the nucleus has been well-documented (56,85). To show that client proteins interact with Hsp90 in ADPKD cells, we conducted pull-down experiments with ErbB2 as a representative Hsp90 client protein that may be relevant to ADPKD progression (64). As shown in Fig. 3B, the endogenous proteins, Hsp90 and ErbB2, co-immunoprecipitated from ADPKD cell lysates.

**Fig. 3.**
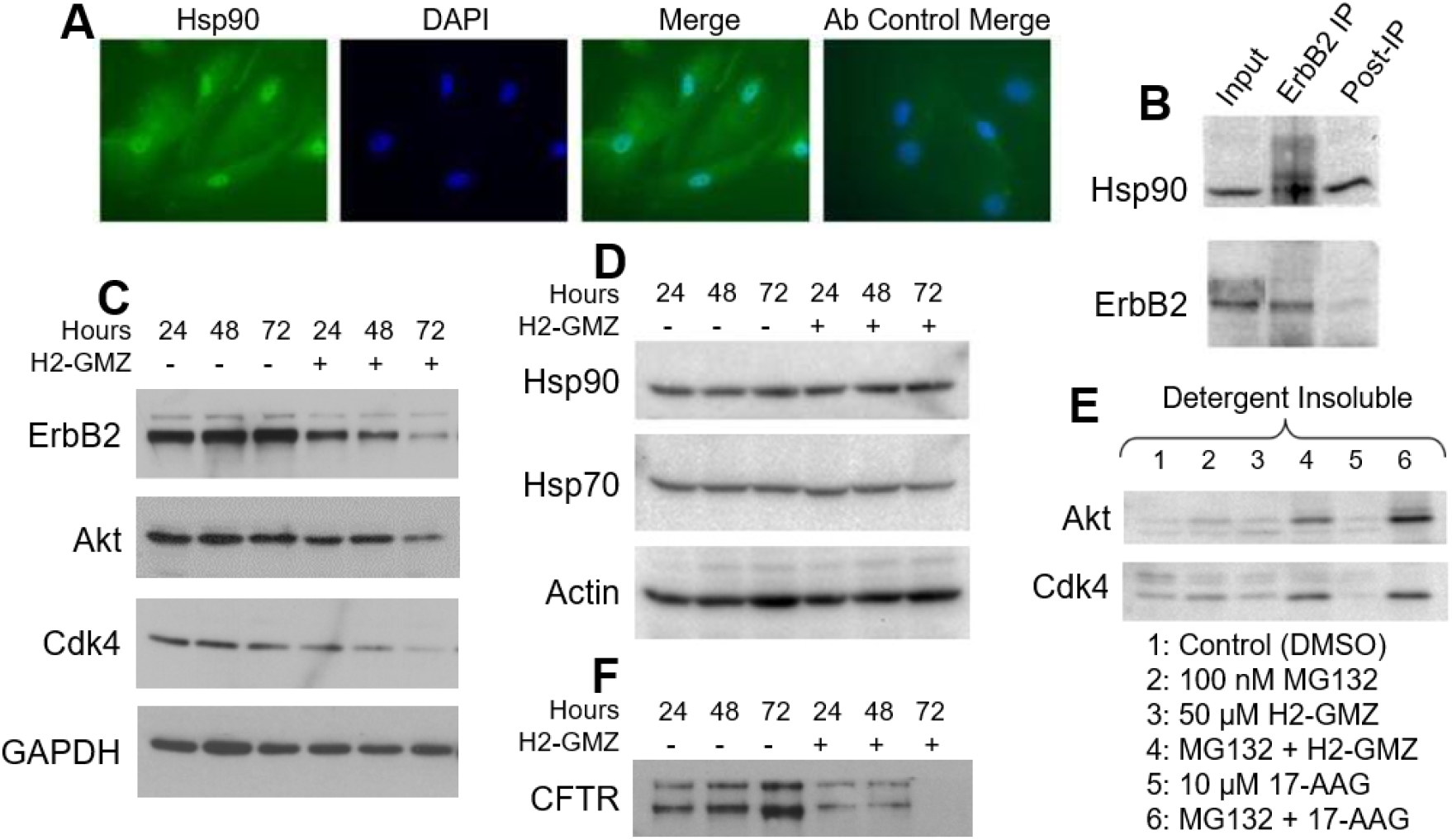
H2-GMZ blocks Hsp90 function in primary human ADPKD cells. **A**, ADPKD cells were stained with anti-Hsp90 antibody (green) to show the presence of Hsp90 in the cytoplasm and nucleus. Cells were counterstained with the nuclear stain DAPI (blue). The control well was incubated with FITC-conjugated secondary antibody alone. **B**, Hsp90 and client protein ErbB2 remain associated in ADPKD cells. ErbB2 was immunoprecipitated from ADPKD cell lysate and the association with Hsp90 was determined by Western blotting. **C** and **D**, H2-GMZ treatment decreases Hsp90 client protein levels. ADPKD cells were treated for 24, 48 or 27 hours with 50 μM H2-GMZ and analyzed by Western blotting. GAPDH and Actin served as internal controls. **E**, H2-GMZ targets Hsp90 client proteins for degradation via the proteasome pathway. ADPKD cells treated with H2-GMZ for 24 hours in the presence or absence of MG132, a proteasome inhibitor, were lysed in buffer containing Triton-X 100 and the insoluble material was pelleted and analyzed by Western blotting. 17-AAG was used as a positive control for the inhibition of the proteasomal pathway. **F**, H2-GMZ treatment decreases CFTR protein levels. All results are representative of three independent experiments with cells from three different ADPKD kidneys.

To determine whether H2-GMZ affects the levels of other Hsp90 pathway proteins, ADPKD cells were treated for various times with 50 μM H2-GMZ and protein levels were determined by Western blotting. H2-GMZ treatment decreased the amounts of the known client proteins ErbB2, Akt and Cdk4, all of which regulate cell proliferation (Fig. 3C). Treatment with H2-GMZ did not increase the levels of Hsp90 or Hsp70 (Fig. 3D). Thus, H2-GMZ appears to affect Hsp90 client proteins without eliciting the heat-shock response, a highly desirable outcome for an Hsp90 pathway inhibitor.

### H2-GMZ treatment leads to degradation of Hsp90 client proteins through the proteasome pathway

Hsp90 inhibitors such as 17-AAG interfere with the chaperone function of Hsp90 (86), thus targeting client proteins to the ubiquitin-mediated proteasomal degradation pathway. Simultaneous inhibition of Hsp90 function and the proteasome pathway would result in the accumulation of ubiquitinated client proteins in the detergent-insoluble fraction of the cell lysate (87). To check whether H2-GMZ targets Hsp90 client proteins for degradation via the proteasome pathway, we treated primary human ADPKD cells with H2-GMZ for 24 h in the presence or absence of the proteasome inhibitor MG132 and determined the levels of Hsp90 client proteins in the detergent (Triton-X 100)-soluble and insoluble fractions from the cell lysate. Fig. 3E shows the accumulation of representative client proteins Akt and Cdk4 in the detergent-insoluble fraction following treatment with H2-GMZ and MG132 (Lane 4). 17-AAG served as positive control (Lane 6).

In our experiments, H2-GMZ appeared to inhibit key signaling proteins that require Hsp90 for maturation or activity. Newly synthesized CFTR protein remains dependent on Hsp90 for functional maturation in the ER, and the Hsp90 inhibitor geldanamycin leads to accelerated degradation of CFTR protein (88). This prompted us to determine the status of CFTR in ADPKD cells following H2-GMZ treatment. We found that H2-GMZ treatment led to a robust inhibition of CFTR protein levels in a time-dependent manner (Fig. 3F). Thus H2-GMZ seems to affect CFTR in multiple ways, as a direct channel inhibitor and by causing a decrease in CFTR protein. As such, H2-GMZ should be an efficient inhibitor of Cl^−^-dependent fluid secretion by ADPKD cyst-lining cells.

### H2-GMZ inhibits ADPKD cell migration

ADPKD cells exhibit significantly higher chemotactic migration in response to EGF compared to normal cells (89). Hence blocking the abnormal motility of ADPKD cells could have a normalizing therapeutic effect. To examine whether H2-GMZ inhibits ADPKD cell motility, we performed a cell migration/wound-healing assay. As little as 10 µM H2-GMZ significantly inhibited the migration of ADPKD cells in 4 h (Fig. 4A, B). The inhibition lasted up to 24 hours and was dose dependent.

**Fig. 4.**
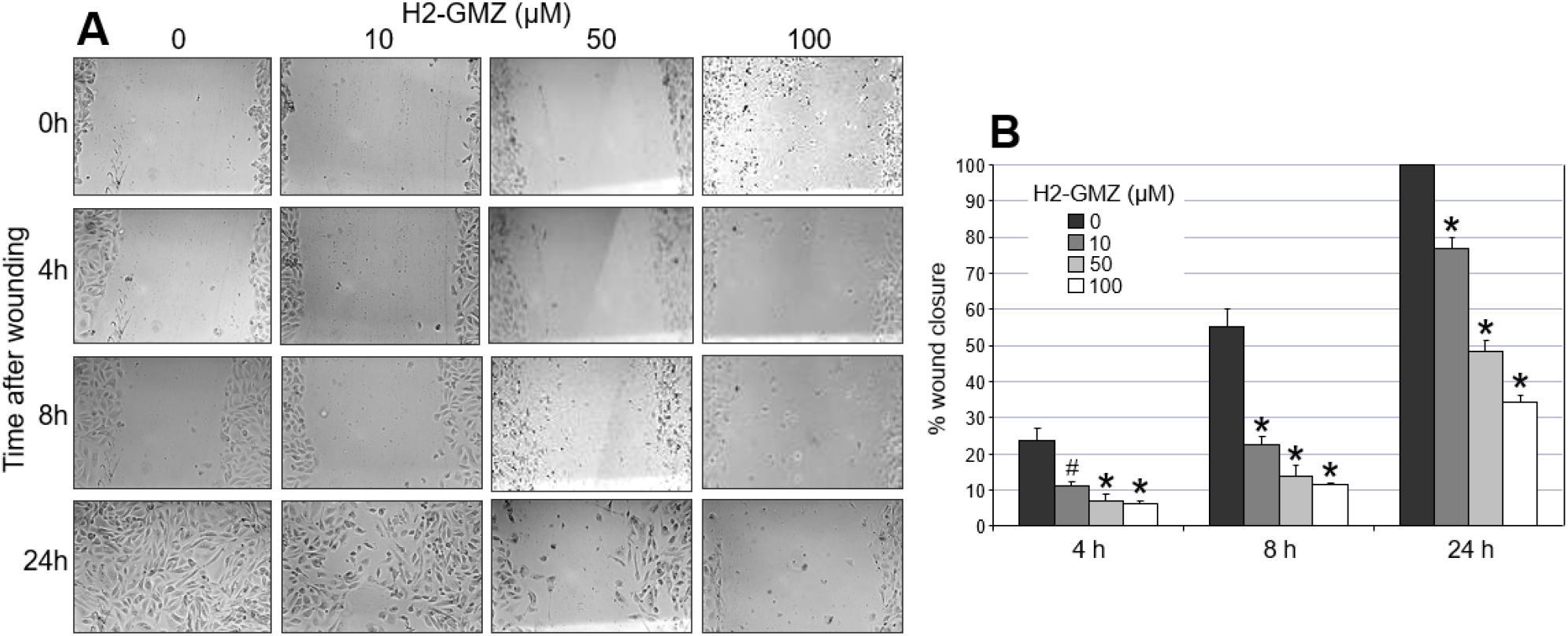
H2-GMZ treatment inhibits cell migration. **A**, Confluent monolayers of human ADPKD cells were scratched with a P20 pipette tip, then treated with increasing concentrations of H2-GMZ. Wound images were captured at 10x magnification immediately after wounding and at 4, 8 and 24 hours post-wounding. **B**, Percentage of wound closure was calculated by subtracting the area of the wound at any given time point from the 0 h wound area expressed as a percentage of the 0 h wound area. Statistical analysis was done to compare the H2-GMZ-treated cells to untreated cells at each time point using One-way ANOVA and the Tukey HSD test for pair-wise comparisons. **#** : p < 0.05, **\***: p < 0.01. The assay was done in triplicate.

Actin bundling is essential for maintaining cell structure, cytoskeletal dynamics and cell motility. In the original screen for proteins binding to GMZ, Tash and colleagues (52) identified the eukaryotic translation elongation factor eEF1A1 as binding strongly to GMZ. eEF1A regulates actin bundling and organization separately from its function in translation (90,91). Since GMZ did not appear to inhibit nucleotide binding by eEF1A1, they hypothesized that GMZ might inhibit actin bundling. To determine whether H2-GMZ can inhibit actin cytoskeletal organization, human ADPKD cells and mouse M-1 cells were stained with phalloidin conjugated with the fluorophore TRITC (Fig. 5A, B). Untreated cells exhibited leading edges (lamellipodia) characteristic of migrating cells, whereas H2-GMZ-treated cells were smaller, without leading edges and with less filamentous actin (F-actin), consistent with an effect on cell motility (Fig. 4B).

**Fig. 5.**
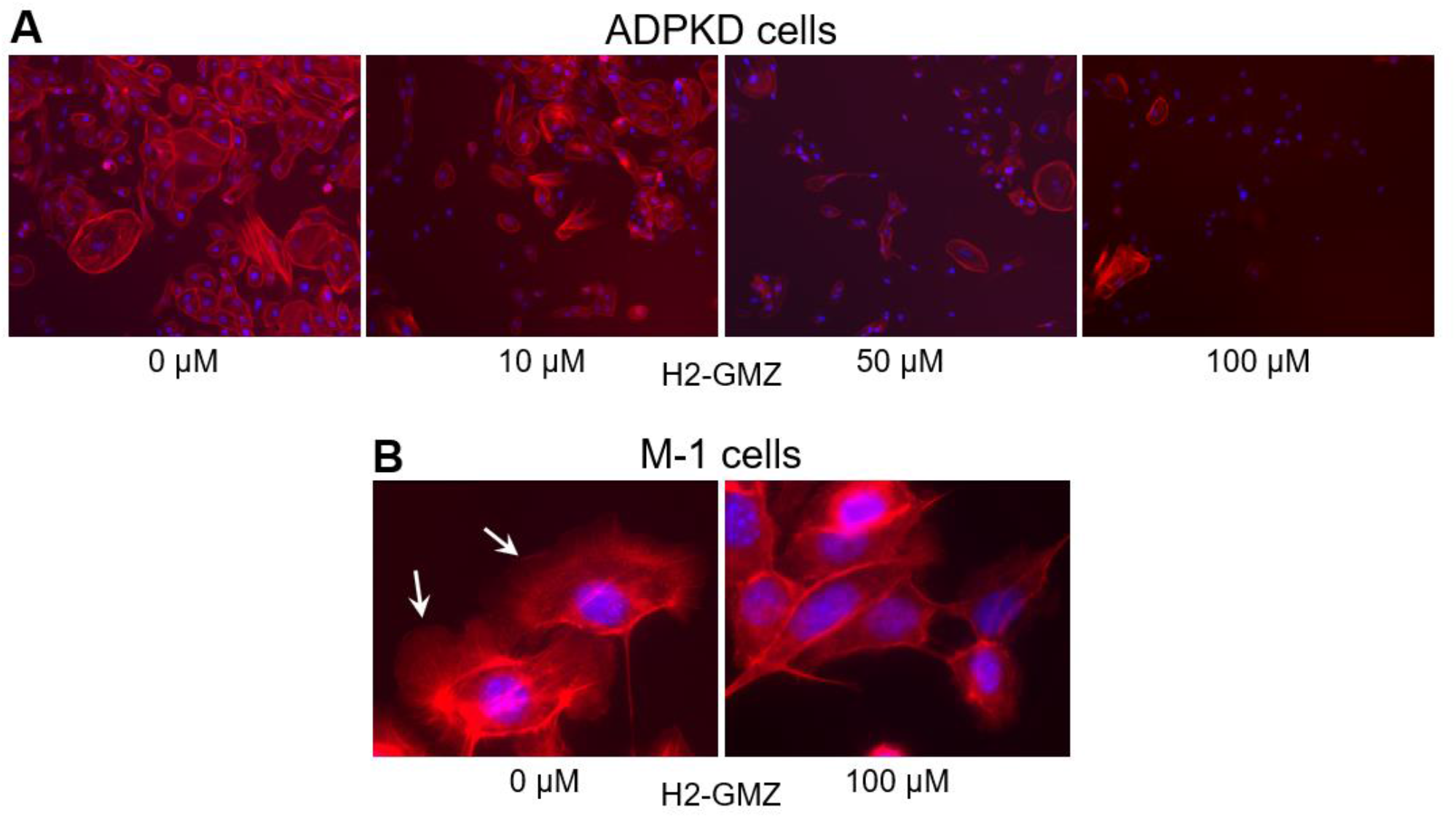
H2-GMZ treatment affects the actin cytoskeleton. **A** and **B**, Human ADPKD cells or mouse M-1 cortical collecting duct cells were treated with increasing concentrations of H2-GMZ and stained with phalloidin. Images were captured at 10x (**A**) or 60x (**B**) using the same exposure settings for treated and untreated cells. Red color shows phalloidin bound to actin filaments in the cell. Blue indicates DAPI-stained nuclei. The assay was done in triplicate.

### H2-GMZ treatment decreases cystic index in metanephric organ culture

We previously showed that CFTR channels are functional in embryonic kidneys and are required for cAMP-driven cyst-like tubule expansion (67). Since H2-GMZ blocked both cell proliferation and CFTR-mediated fluid secretion in ADPKD cells, we examined the effectiveness of H2-GMZ in inhibiting cAMP-dependent cyst growth and enlargement in mouse embryonic kidneys in organ culture. For this, we used kidneys from *Pkd1*^*m1Bei*^ +/− or −/− embryos (81). As shown in Figure 6, 5 μM H2-GMZ significantly inhibited cyst formation.

**Fig. 6.**
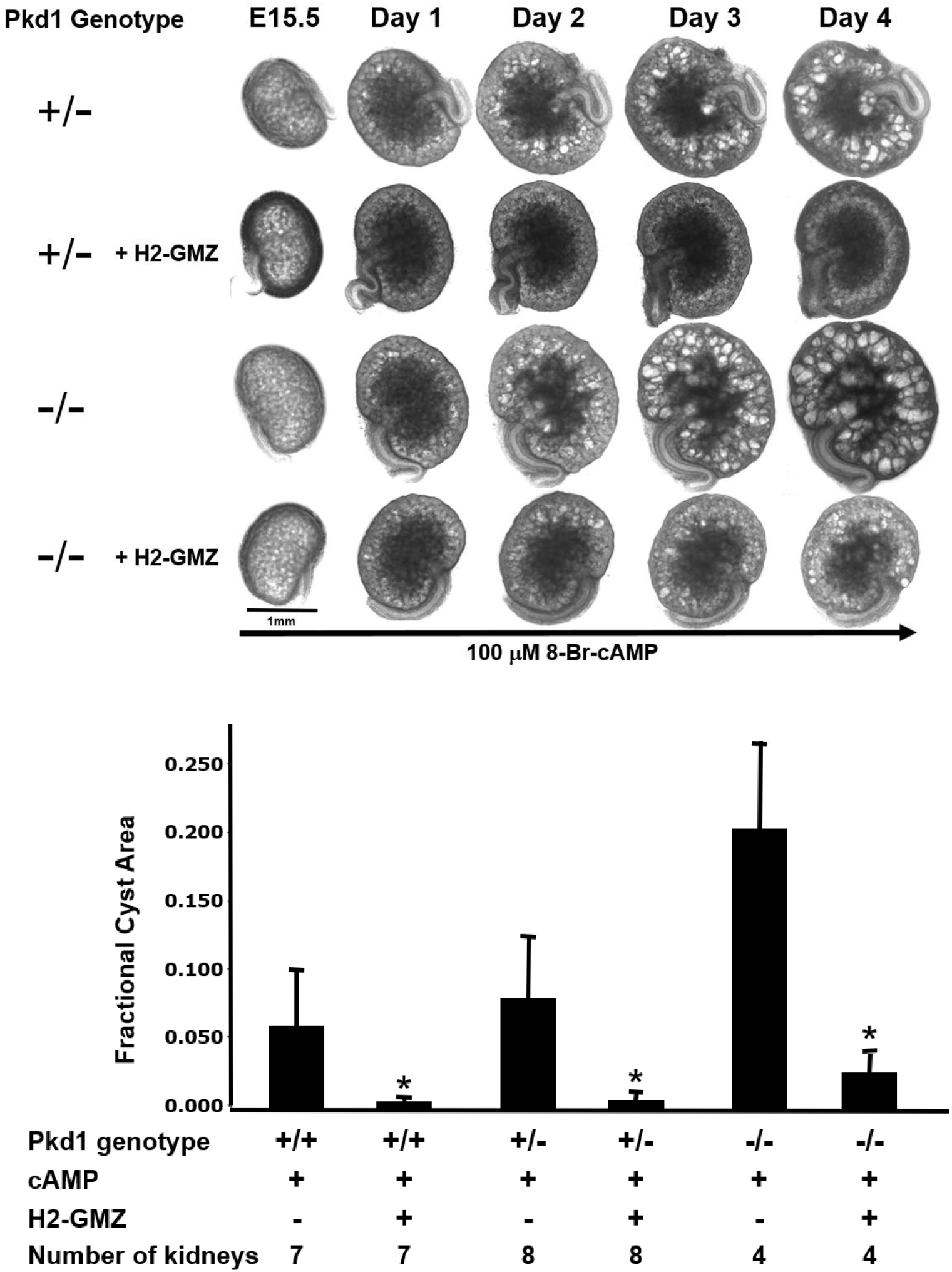
H2-GMZ treatment reduces the cyst burden in cAMP-treated metanephric kidneys. Embryonic day15.5 mouse kidneys from Pkd1 +/− and −/− mice were plated on Transwell membranes and treated with 100 μM cAMP with or without 5 μM H2-GMZ for four days. H2-GMZ treatment reduced the cystic index of treated kidneys. Fractional cyst area is the total area of all the cysts per kidney represented as a fraction of the total area of the kidney. The number of kidneys of each genotype is listed.**\***, the effect was statistically significant at p < 0.01 as determined by Student’s t-test.

H2-GMZ was also effective when it was added a day after cyst initiation (Supplemental Fig. 2) and was partially effective when it was washed out of the medium following a 24 h treatment period (Supplemental Fig. 3). LND also inhibited cyst growth in *Pkd1*^*m1Bei*^ +/− metanephric kidneys (Supplemental Fig. 4) but did not appear to be as effective as the same dose of H2-GMZ. Five μM LND did not inhibit cyst growth in *Pkd1*^*m1Bei*^ −/− metanephric kidneys (data not shown).

Next, we stained H2-GMZ treated metanephric kidneys for PCNA, a marker for cell proliferation. There was a marked decrease in the intensity of PCNA staining in treated kidneys compared to controls (Supplemental Fig. 5A, B). As expected for growing embryonic kidneys, there were many brightly stained vesicles and S-shaped bodies visible in the untreated kidney while there were very few in the H2-GMZ treated kidney.

### H2-GMZ treatment decreases cyst progression *in vivo* in a *Pkd1* mouse model

To determine the effects of H2-GMZ *in vivo*, treatment was carried out on *Pkd1*^*flox/flox*^: *Pkhd1-Cre* mice using daily i.p. injections of 20 mg/kg H2-GMZ from postnatal day 8 (PN8) to 18 (PN18). Mice treated with H2-GMZ had significantly smaller kidneys and increased renal parenchyma (Fig. 7A), reduced cystic index (Fig. 7B), decreased kidney weight to body weight (KW/BW), and improved blood urea nitrogen (BUN) levels (Fig. 7B). Survival studies (Fig. 7C) showed significantly longer survival for H2-GMZ-treated mice (Control vs H2-GMZ: 28.8 ± 5 vs 67.8 ± 23; p < 0.01) indicating that H2-GMZ may be effective in prolonging renal function.

**Fig. 7.**
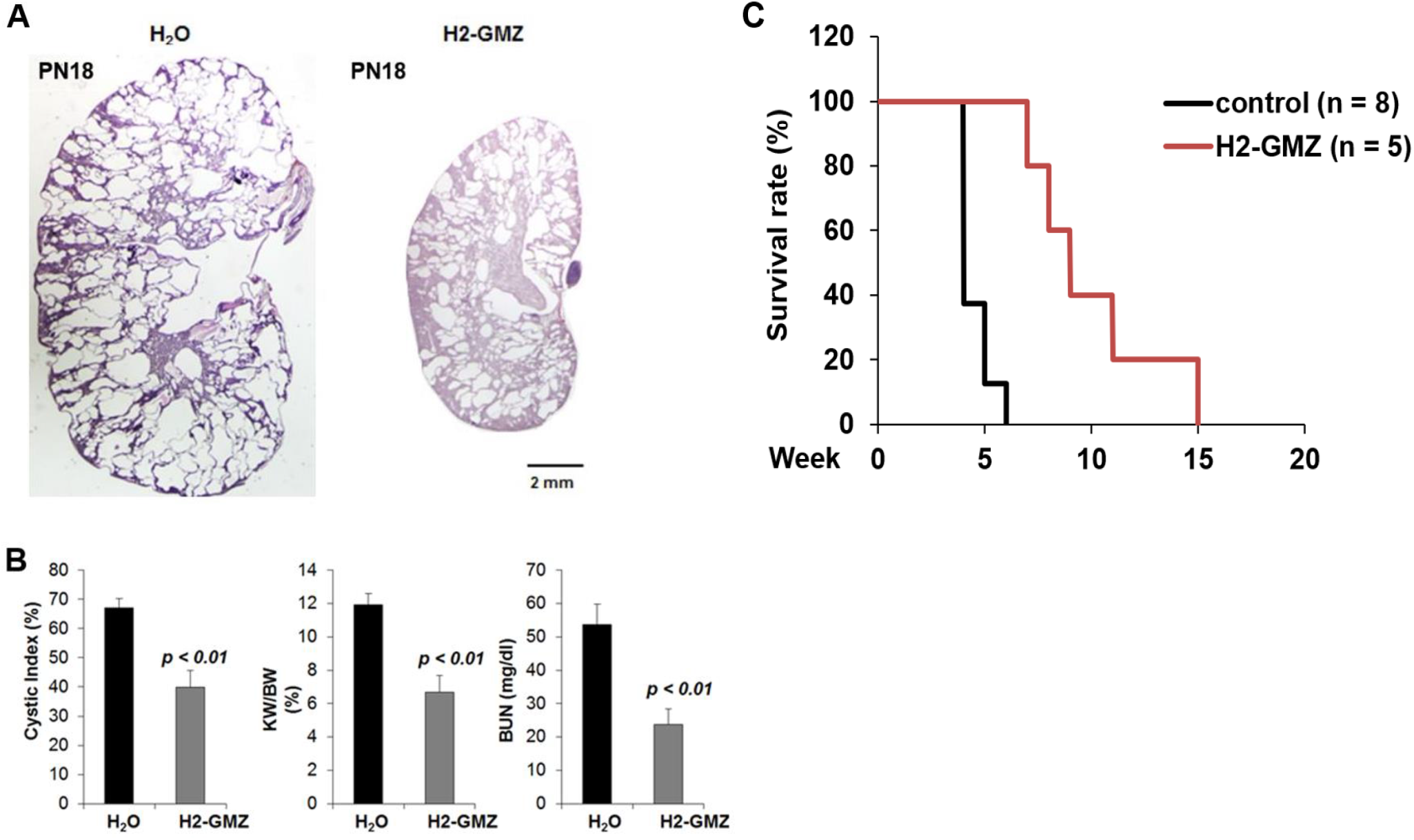
H2-GMZ treatment decreases cystic burden in vivo in a Pkd1 mouse model. H2-GMZ treatment was carried out on Pkd1^flox/flox^:Pkhd1-Cre mice using daily i.p. injections of 20 mg/kg H2-GMZ from postnatal day 8 to 18. **A** and **B**, Mice treated with H2-GMZ had a significantly reduced cystic index, smaller kidneys and increased renal parenchyma, decreased kidney weight to body weight (KW/BW) and improved blood urea nitrogen (BUN) levels. **C**, Survival studies showed significantly longer survival for H2-GMZ-treated mice (Control vs H2-GMZ: 28.8±5 vs 67.8±23; p < 0.01).

## DISCUSSION

In this study, we investigated the potential of a new class of drugs for ADPKD. We provide evidence that the novel indazole carboxylic acid derivative, H2-GMZ, is effective in inhibiting fluid secretion and cell proliferation in human ADPKD cells, mouse metanephric kidneys and in a rapidly progressive mouse model of PKD. To further understand the mechanism of action of H2-GMZ, we explored its effects on the activities of CFTR, Hsp90, and eEF1A1, three proteins through which H2-GMZ appears to mediate its actions in the cell. Human ADPKD cell monolayers responded to H2-GMZ treatment with decreased CFTR-mediated anion current. We also found that H2-GMZ decreased the levels of a number of Hsp90 client proteins, including CFTR, and targeted them for degradation through the ubiquitin-proteasome pathway. Many of these Hsp90 client proteins are required for cell growth and proliferation. H2-GMZ also inhibited the motility of ADPKD cells and decreased F-actin levels, consistent with an inhibition of the actin-bundling properties of eEF1A1. Furthermore, low micromolar concentrations of H2-GMZ were effective in inhibiting cyst formation in a *Pkd1* mutant mouse metanephric organ culture model in which a number of processes mediate the growth and enlargement of cysts in response to cAMP (67). Finally, daily injection of H2-GMZ in a postnatal *Pkd1* mouse model showed inhibition of cystogenesis and increased survival indicating that H2-GMZ effectively targets the kidney *in vivo*.

Hsp90 function is critical for protein folding and function in eukaryotic cells. Protein molecules exist in a crowded state in the cell, which favors inter-molecular interactions but risks misfolding and aggregation (56,57). Hsp90 uses energy derived from ATP to promote the protein conformation required for proper functioning. In various neoplastic disorders, Hsp90 and its client proteins are overexpressed and/or hyperactive (92). This leads to increased signaling and uncontrolled cell growth and division. Targeting Hsp90 to reduce its activity has proven to be beneficial in curbing the growth of many tumors (61,63,86,92-94). The Hsp90 inhibitors STA-2842 and STA-9090 (ganetespib) have been shown to be effective in reducing cyst size and disease progression in mouse models PKD (75-77). As ADPKD has many of the characteristics of a neoplastic disorder (95), we believe this approach may be useful in treating patients with ADPKD.

In ADPKD, renal epithelial cysts grow and enlarge through a process of abnormal cell proliferation and increased fluid secretion. Each cyst could be considered a fluid-filled tumor (96). Apical Cl^−^ transport by CFTR drives net fluid secretion into the cyst lumen in response to cAMP (12,16,97,98). Cyst-lining cells are poorly differentiated, and express elevated proto-oncogenes (99). Overexpression of oncogenes c-myc, rasT24, and SV40 large T antigen leads to PKD in transgenic mouse models (100-102). A key pathway that is abnormally activated in PKD is the MEK-ERK pathway, as seen also in many types of cancer (13,19,71-73,103,104). In addition, there is evidence for activation of the EGF family receptors, IGF receptors and β-catenin in many types of cancer as well as PKD (64,74,105,106). A number of these pathways need Hsp90 for sustained activation. There may be higher than normal levels of certain proteins in ADPKD cells that would utilize Hsp90 for their continued aberrant function. For example, the EGFR family receptor ErbB2 (Her2), an Hsp90 client protein, is overexpressed in PKD (64,65). Cyst-lining renal epithelial cells also overexpress CFTR (97,98,107). For these reasons we expect that an inhibitor of the Hsp90 pathway would preferentially target the cystic epithelium rather than the normal renal parenchyma.

H2-GMZ is a novel derivative of the anti-cancer drug lonidamine (LND), used to treat various types of cancer (47,108-110). LND is known to inhibit cellular glucose metabolism, possibly by its effect on mitochondrial hexokinase (110,111). Drug resistant breast cancer cells and malignant glioma cells were found to be especially sensitive to LND (110,111). In addition, LND inhibits CFTR activity in cultured epididymal cells, presumably by direct binding to the pore of the channel (48-50). While the exact mechanism of anti-tumor activity of LND is unknown, it is most likely a combination of the activities mentioned above and possibly additional activities. LND was originally described as an anti-spermatogenic drug (112). In an effort to develop better male contraceptives, various analogs of LND were synthesized among which GMZ showed the greatest potency and the least side effects (52,55). Since PKD is a chronic condition that might require a lifetime of therapy, a drug with excellent safety and toxicity profiles would be the ideal candidate for PKD therapy. In animal studies, GMZ was very well tolerated (55).

In studies to understand the mechanism of action of GMZ, it was observed that GMZ affects two proteins: the constitutive isoform of Hsp90, Hsp90AB1, and the eukaryotic elongation factor eEF1A1 (52). The mechanism by which GMZ inhibits Hsp90 is currently being investigated. Neither geldanamycin (which binds to the N-terminus of Hsp90) nor a novobiocin derivative (which binds the C-terminus) could compete with GMZ for Hsp90 binding (52). Therefore, it is possible that GMZ binds to a different site on Hsp90 or that it binds one of the co-chaperones in the Hsp90 super complex. In fact, subsequent studies have demonstrated that the Hsp90-cdc37 chaperone complex was disrupted following H2-GMZ treatment in the ErbB2-overexpressing cancer cell line SKBr3 (53). Hsp90 inhibition by GMZ was also confirmed by a decrease in client proteins ErbB2 and Akt (52,53).

In testing the effect of H2-GMZ on Hsp90-mediated activation of key signaling proteins, we found that H2-GMZ treatment effectively decreased ErbB2 and Akt. ErbB2 (Her2) is known to be upregulated in ADPKD cells compared to normal renal epithelial cells and treatment with ErbB2 inhibitors was found to slow cyst growth in a mouse model of ADPKD (64). ADPKD cyst-lining epithelial cells exhibit misregulation of the PI3K-Akt-mTOR pathway (69,113). The cyclin-dependent kinase Cdk4 is an Hsp90 client protein which phosphorylates retinoblastoma protein (Rb) during the G1 phase of the cell cycle. The H2-GMZ mediated decrease in Cdk4 and hyperphosphorylated Rb could contribute to the observed decrease in cell proliferation. Consistent with H2-GMZ being an Hsp90 inhibitor, we showed that the unfolded client proteins were directed to the proteasome pathway for degradation.

One of the Hsp90 client proteins, the Cl^−^ channel CFTR, plays an important role in the pathogenesis of ADPKD by mediating fluid secretion into the cyst lumen (67,114), and CFTR inhibitors have proven effective in reducing cyst growth (66). We found that CFTR secretion decreases rapidly following treatment with H2-GMZ. Furthermore, CFTR protein levels decrease markedly 24 to 72 hours after H2-GMZ treatment. Since LND directly binds to and blocks CFTR, it is possible that H2-GMZ acts similarly. Thus H2-GMZ could be inhibiting CFTR channel activity via direct binding and also indirectly through Hsp90, making this drug an especially attractive candidate for ADPKD therapy.

ADPKD cell motility and migration could be involved in cyst formation and PKD progression since ADPKD cells as a monolayer on solid support migrate faster than normal cells to achieve wound closure (89). The integrity of the actin cytoskeleton is critically important to the ability of cells to migrate. It is known that LND significantly inhibits endothelial cell migration and invasiveness (115). LND treatment caused the disappearance of actin stress fibers and rearrangement of intermediate filaments and microtubules (116). We found that H2-GMZ treatment effectively blocked ADPKD cell migration, perhaps by its effect on eEF1A1 (52). eEF1A1 is essential for regulation of the actin cytoskeleton (91,117). Previous experiments showed that GMZ does not affect the GDP/GTP nucleotide binding properties of eEF1A1, which are required for it to mediate elongation during translation (52). Therefore it was postulated that GMZ might inhibit the bundling of F-actin by eEF1A1, a function separate from its role in protein translation (91). Our results indicate that H2-GMZ might disrupt cell migration by inhibiting actin polymerization. It is also possible that H2-GMZ could mediate its effect on cell migration independent of eEF1A1.

A drug like H2-GMZ with low toxicity and multiple cellular targets would be ideal for PKD treatment. One of the drawbacks of traditional Hsp90 inhibitors such as geldanamycin derivatives is the compensatory induction of Hsp90 and Hsp70 (118-120) thus reducing their effectiveness over time. In our studies, H2-GMZ did not induce Hsp90 or Hsp70 even after 72 hours of treatment. This lack of a heat shock response in treated cells could make the drug more effective in targeting the Hsp90 machinery. While H2-GMZ was explored as a potential male contraceptive, its anti-spermatogenic effect may not be considered a drawback for every male patient, or for female patients. Depending on the stage of the disease and the age and health of the patient, the long-term advantages may outweigh this potential disadvantage, thus justifying the use of H2-GMZ or other lonidamine derivatives in the treatment of ADPKD, although an effort to develop H2-GMZ analogs without male reproductive effects would be a worthwhile goal of future studies. Ultimately, the ideal treatment for PKD may be a combination of therapies in low tolerable doses that hit multiple critical targets including both cell proliferation and fluid secretion to slow cyst growth and enlargement and to preserve kidney function over the long term.

## ACKNOWLEDGEMENTS

Supported by grants from the KU Institute for Advancing Medical Innovation (IAMI), NIH (P30 DK106912) and PKD Foundation to JPC, NIH (R01 DK081579) to DPW, and the PKD Biomarkers, Biomaterials, and Cellular Models Core of the Kansas PKD Research and Translation Core Center (U54 DK126126). The authors thank Dr. Peter Igarashi and Dr. Stefan Somlo for the *Pkd1*^*flox/flox*^: *Pkhd1-Cre* mice.

