## Supplemental Figures for "The Lonidamine Derivative H2-Gamendazole Reduces Cyst Formation in Polycystic Kidney Disease"

### Supplemental Fig. 1

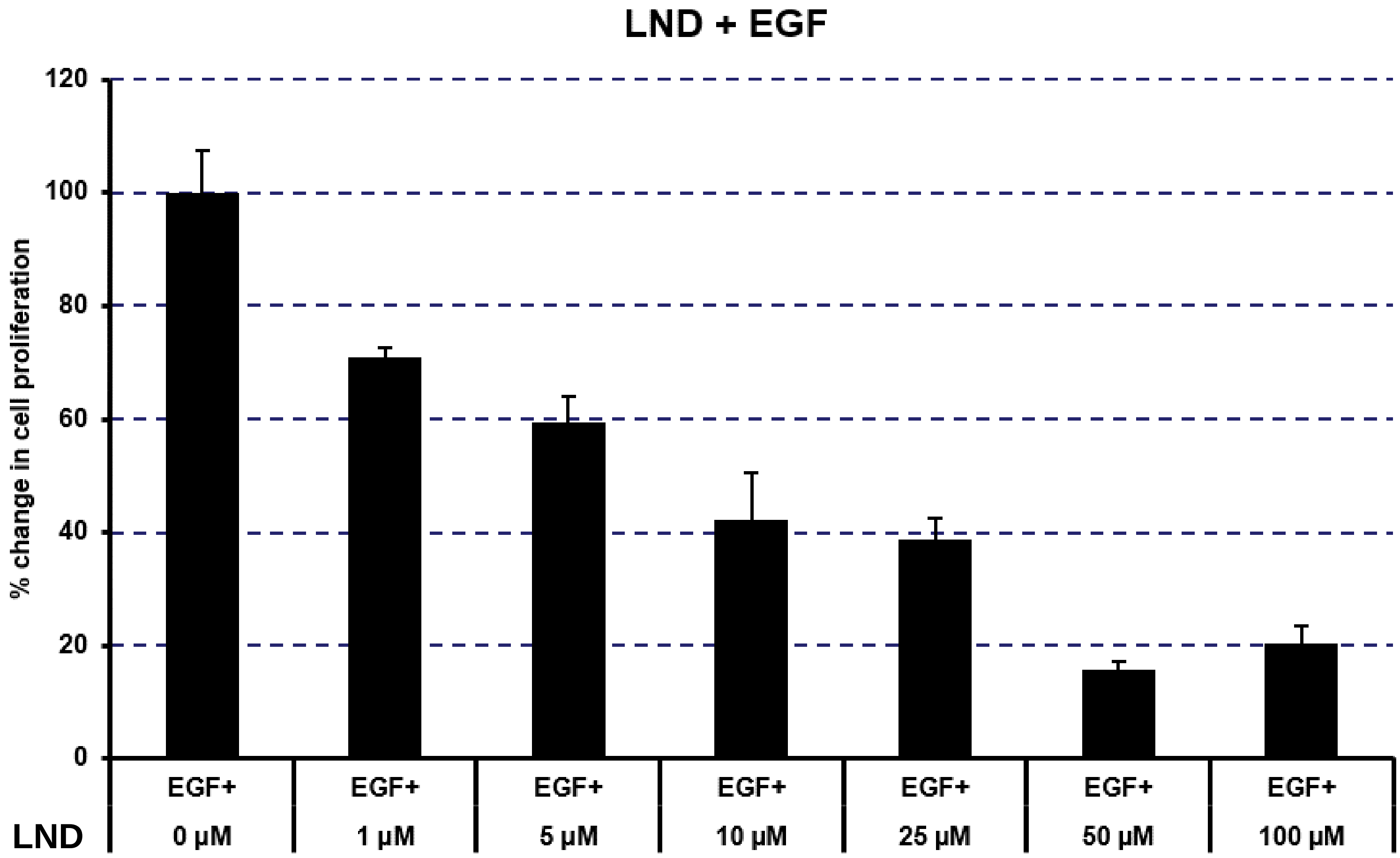


**Supplemental Fig. 1. Dose-dependent decrease in the proliferation of primary human ADPKD cells treated with LND.** Dose response of ADPKD cells stimulated with 25 ng/ml EGF and treated with the indicated doses of LND. After 72 hours of treatment, MTT assays were performed. In each group, results were expressed as a percentage of the control group (not treated). The error bars indicate a standard error.

### Supplemental Fig. 2

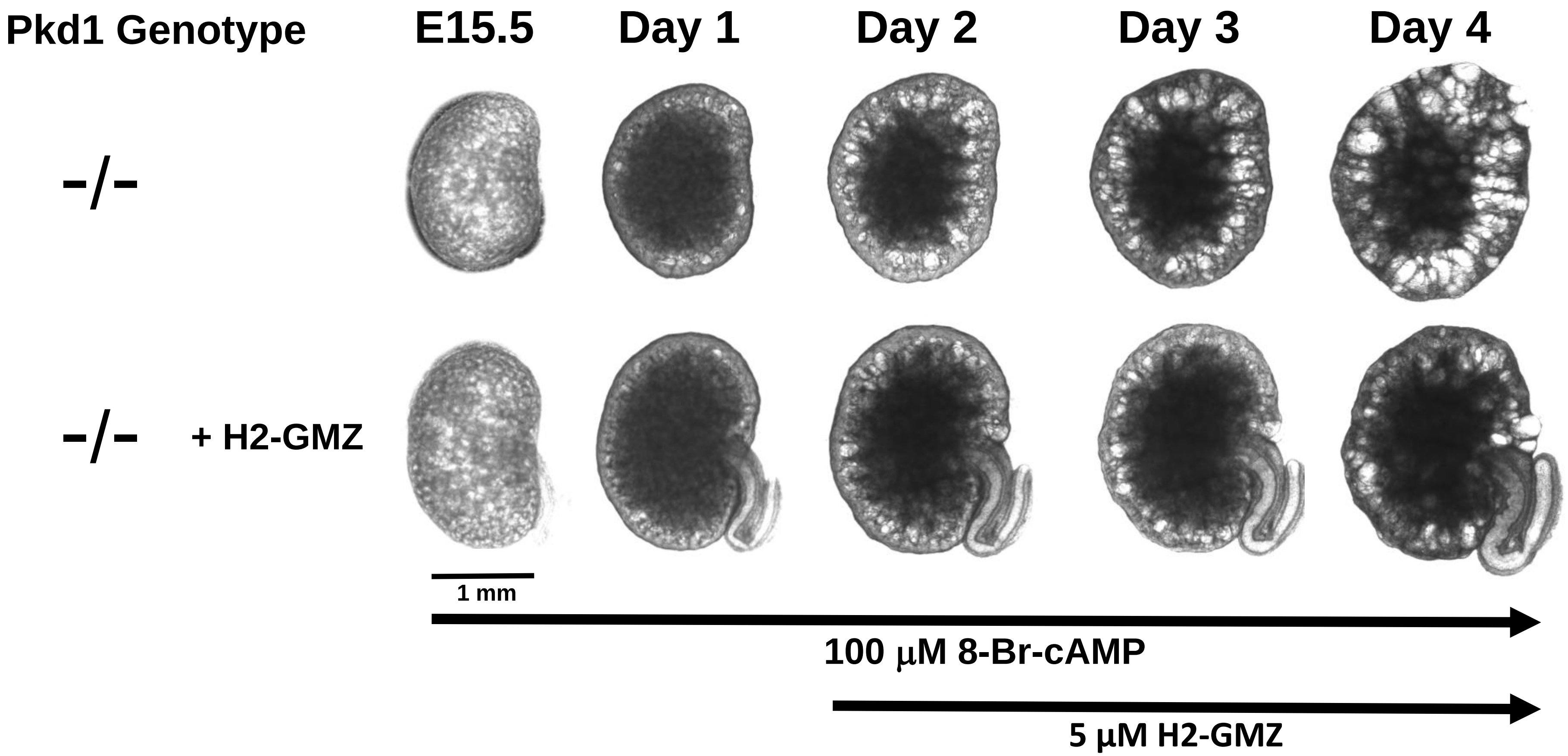


**Supplemental Fig. 2. Delayed addition of H2-GMZ is effective in decreasing cyst growth.** Addition of 5 μM H2-GMZ on Day 2 Pkd1 -/- metanephric kidneys was effective in reducing cystic dilation.

Supplemental Fig. 3

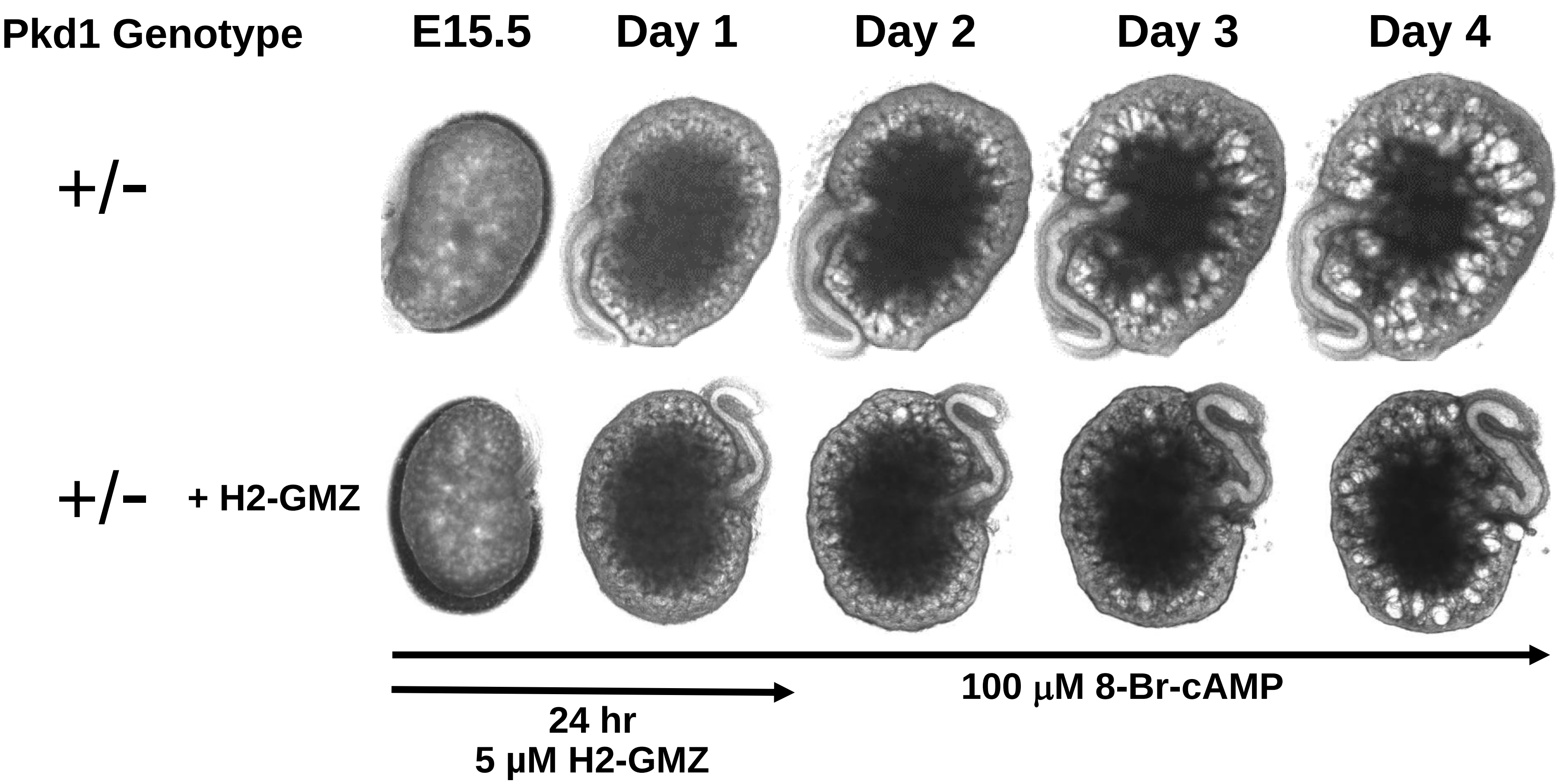


**Supplemental Fig. 3. The inhibitory effects of H2-GMZ with a 24-hour treatment.** Addition of 5 μM H2-GMZ to Pkd1 +/- kidneys was terminated after 24 hours and showed partial effectiveness.

#### Supplemental Fig. 4

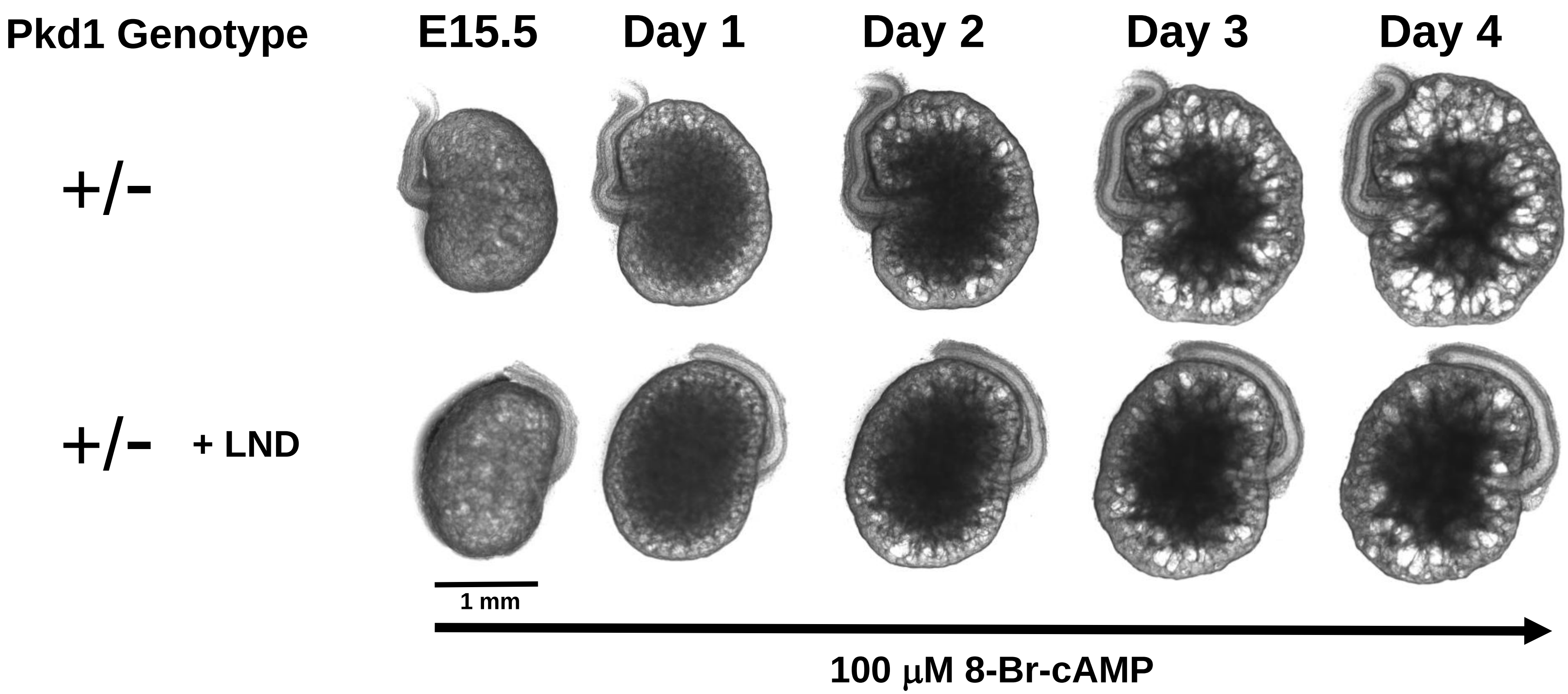


**Supplemental Fig. 4. LND treatment reduces the cystic dilations in cAMP-treated *Pkd1* +/- metanephric kidneys.** Embryonic day15.5 mouse kidneys from Pkd1 +/- mice were plated on Transwell membranes and treated with 100 μM cAMP with or without 5 μM LND for four days. LND treatment decreased cyst formation in metanephric kidneys.

**Supplemental Fig. 5A**

A

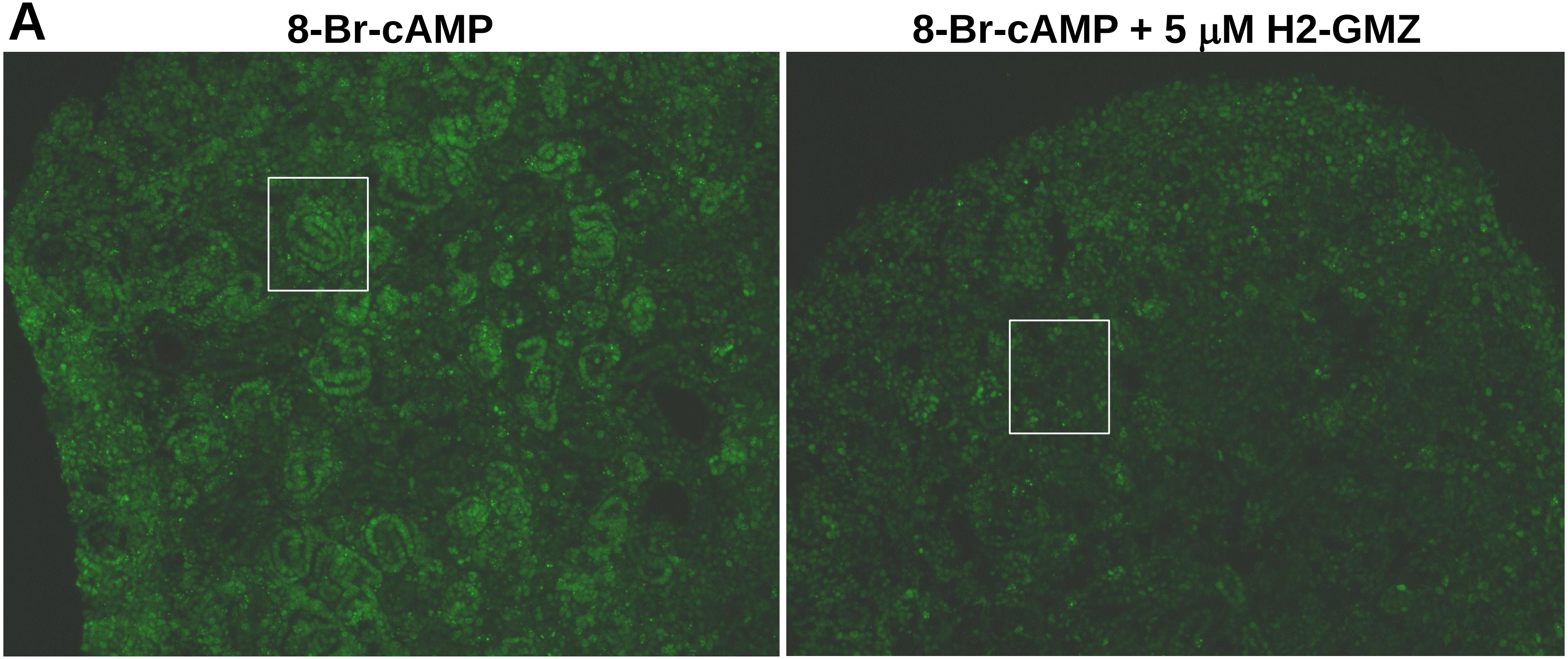


**Supplemental Fig. 5. H2-GMZ inhibits proliferation in metanephric kidneys as determined by PCNA staining. A,** E15.5 wild type metanephric kidneys were treated with 100 μM 8-Br-cAMP and 5 μM H2-GMZ for four days. Frozen sections were stained for PCNA using double immunofluorescence. The secondary antibody was conjugated to FITC (green). **B**, Higher magnification of boxed areas showing brighter PCNA staining in the tubule cells of the control kidney.

#### Supplemental Fig. 5B

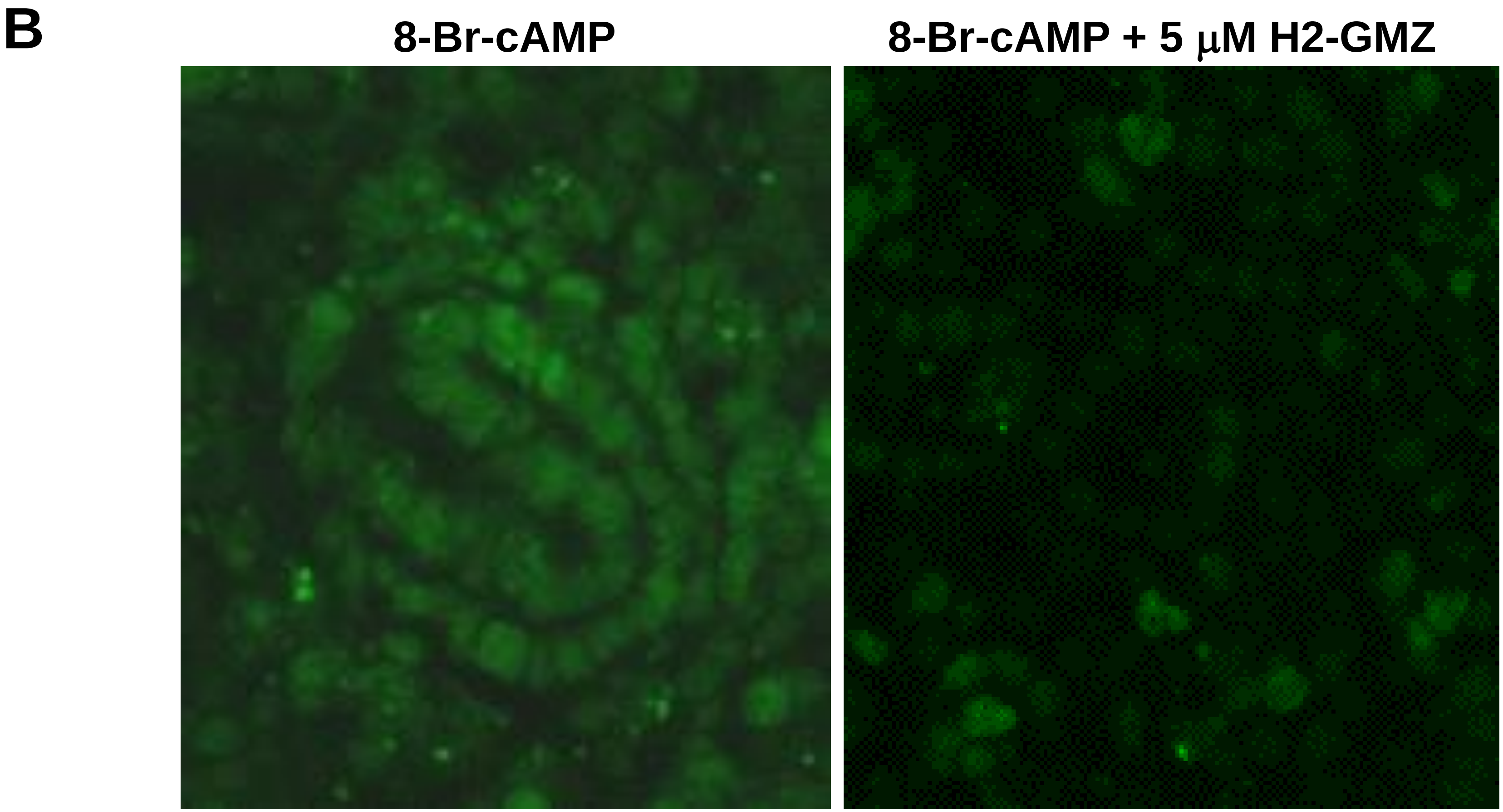


**Supplemental Fig. 5. H2-GMZ inhibits proliferation in metanephric kidneys as determined by PCNA staining. A,** E15.5 wild type metanephric kidneys were treated with 100 μM 8-Br-cAMP and 5 μM H2-GMZ for four days. Frozen sections were stained for PCNA using double immunofluorescence. The secondary antibody was conjugated to FITC (green). **B**, Higher magnification of boxed areas showing brighter PCNA staining in the tubule cells of the control kidney.
